# BOLD Decoding of Individual Pain Anticipation Biases During Uncertainty

**DOI:** 10.1101/675645

**Authors:** Molly Kadlec, Duygu Tosun, Irina Strigo

**Author notes:** Corresponding Author: Irina Strigo, Ph.D., Emotion and Pain Laboratory, San Francisco Veterans Affairs Health Care Center, 4150 Clement Street, San Francisco, CA 94121.

## Abstract

A prominent model of pain as a predictive cue posits that anticipation shapes pain transmission and ultimately pain experience. Consistent with this model, the neural mechanisms underlying pain anticipation have the power to modulate pain experience thus understanding pain predictions, particularly during uncertainty, may allow us to ascertain measures indicative of intrinsic anticipation biases. Understanding such biases moves way to precision pain management, as it can guide the individualized treatment. To examine individual pain anticipation biases, we applied machine-learning-based neural decoding to functional magnetic resonance imaging (fMRI) data acquired during a pain-anticipation paradigm to identify individualized neural activation patterns differentiating two certain anticipatory conditions, which we then used to decode that individual’s uncertain anticipatory condition. We showed that neural patterns representative of the individualized response during certain anticipatory conditions were differentiable with high accuracy and, across individuals, most commonly involved neural activation patterns within anterior short gyrus of the insula and the nucleus accumbens. Using unsupervised clustering of individualized decodings of anticipatory responses during uncertain conditions, we identified three distinct response profiles representing subjects who, in uncertain situations, consistently anticipated high-pain (i.e., negative bias), subjects who consistently anticipated low-pain (i.e., positive bias), and subjects whose decoded anticipation responses were depended on the intensity of the preceding pain stimulus. The individualized decoded pain anticipation biases during uncertainty were independent of existence or type of diagnosed psychopathology, were stable over one year timespan and were related to underlying insula anatomy. Our results suggest that anticipation behaviors may be intrinsic, stable, and specific to each individual. Understanding individual differences in the neurobiology of pain anticipation has the potential to greatly improve the clinical pain management.

## Introduction

Anticipation, an inherent component of emotional processing, shapes how humans respond to both rewarding and aversive stimuli. In the context of pain, how an individual anticipates the upcoming noxious stimuli modulates [1–4] how pain is experienced by engaging appropriate pain relief mechanisms [5,6]. Previous studies have shown that by changing an individual’s expectation or attentiveness, their perception of the pain is changed [3,4], as well as their brain response [7–13]. Positive expectancy cues (i.e., the expectation of analgesia) have been shown to reduce pain and result in placebo analgesia, while negative expectancy cues (i.e., expectation of worsening pain) can lead to an increase in reported pain and result in nocebo analgesia [4, 14–18]. Intrinsic differences in an individual’s anticipatory biases could be probed to identify and objectively classify individuals as adaptive, i.e., those with positive bias, who successfully regulate emotional responses to unpleasant stimuli [19, 20], or maladaptive responders, i.e., those with negative bias, who show heightened reactivity, behavioral and cognitive avoidance, and increased threat attention, among others [21–25]. The intrinsic differences in individuals’ anticipatory biases are particularly important when anticipatory cues become ambiguous, since real-life scenarios are often fraught with uncertainty. It has been suggested that uncertainty is a crucial aspect of decision-making as it can lead to prediction errors (incorrectly anticipating a given outcome) [13]. These errors allow individuals to reevaluate the cost-benefit relationship in a given scenario to inform future decisions [13]. In the motivation-decision model of pain, expectation effects are seen through decision-making processes mediated by top-down neural circuitries which either enhance or inhibit nociceptive transmission to drive behavioral responses [13]. This model emphasizes the effects anticipation can have on pain perception, and points to the presence of individual variability in how pain is anticipated as a necessary decision.

Here we assess to what extent individuals’ own neural activity patterns during pain anticipation could identify their positive or negative anticipatory response biases in ambiguous and uncertain situations. Using novel machine learning techniques and single-subject data analytics, our primary aims were to create a subject-specific voxel activation pattern distinguishing two levels of cued pain anticipation (i.e., anticipating “high” pain stimulus versus anticipating “low” pain stimulus) and use it to decode that individual’s intrinsic anticipation response patterns during uncertainty. Our secondary aim was to discover whether anticipation response patterns during uncertainty were influenced by demographics, psychopathology, or brain structure. To date, this is the first known study that used a single-subject machine learning approach to distinguish between high and low-pain anticipation patterns in a large heterogeneous cohort of men and women with and without psychopathology.

## Results

### High Accuracy of Single-Subject LASSO Model to distinguish Low and High-pain Anticipation Neural Response Pattern

We conducted functional magnetic resonance imaging (fMRI) in one hundred and forty-seven subjects (50 females; 28 ± 6.8 years old) while subjects performed a pain-anticipation paradigm including two anticipation conditions of certain pain intensity (visual cue indicating whether a high or low-pain stimulus will be delivered), and one anticipation condition of uncertain pain intensity (ambiguous cue indicating either high or low-pain could be experienced). Fifty-seven subjects (22 females) were healthy controls with no current or past history of mental illness or trauma, and ninety (28 females) were part of a “mixed psychiatric” test group (see Methods for details). Each subject received the pain-anticipation paradigm sequence (or run) twice with a randomized order of stimulation conditions within each run (see Methods). Between two pain-anticipation paradigm sequences, each participants performed a total of seven low-pain anticipation conditions, seven high-pain anticipation conditions, and fourteen uncertain pain anticipation conditions. We first examined whether neural patterns representative of the anticipation of high-pain and low-pain were reliably distinguishable in every subject. Due to statistical power limitations of single-subject data analytics, we limited our inquiry (“feature selection”) to twenty-six brain regions, spanning across the insula, anterior cingulate, amygdalae, and dorsal and ventral striatum bilaterally, known to play prominent roles in pain prediction, processing, and relief [13, 26, 27]. The study flow is summarized in **Figure 1**. Using logistic regression analysis with least absolute shrinkage and selection operator (LASSO) regularization, the performance of individualized neural patterns in separating anticipation of high-pain versus anticipation of low-pain was 97.4 ± 7.9% accurate with 98.4 ± 5.9% sensitivity, 96.5 ± 13.1% specificity, and 97.4% area under the curve (AUC) across all subjects, as indicated by the receiver operating characteristic (ROC) curves in **Figure 2**. High-pain and low-pain anticipatory neural patterns were separable in 145 of 147 subjects in our study based on a single-subject accuracy threshold of 75% or greater. Across all subjects’ LASSO models, insular regions were the most frequently (96% of the subjects) included regional neural activity predictors of low-pain versus high-pain anticipation neural pattern separation, with anterior short gyrus being the most frequent (63%). Other highly contributing regions were nucleus accumbens (64%), substantia nigra (61%) and amygdala (60%) (see Supplementary Material 6 for details).

**Figure 1:**
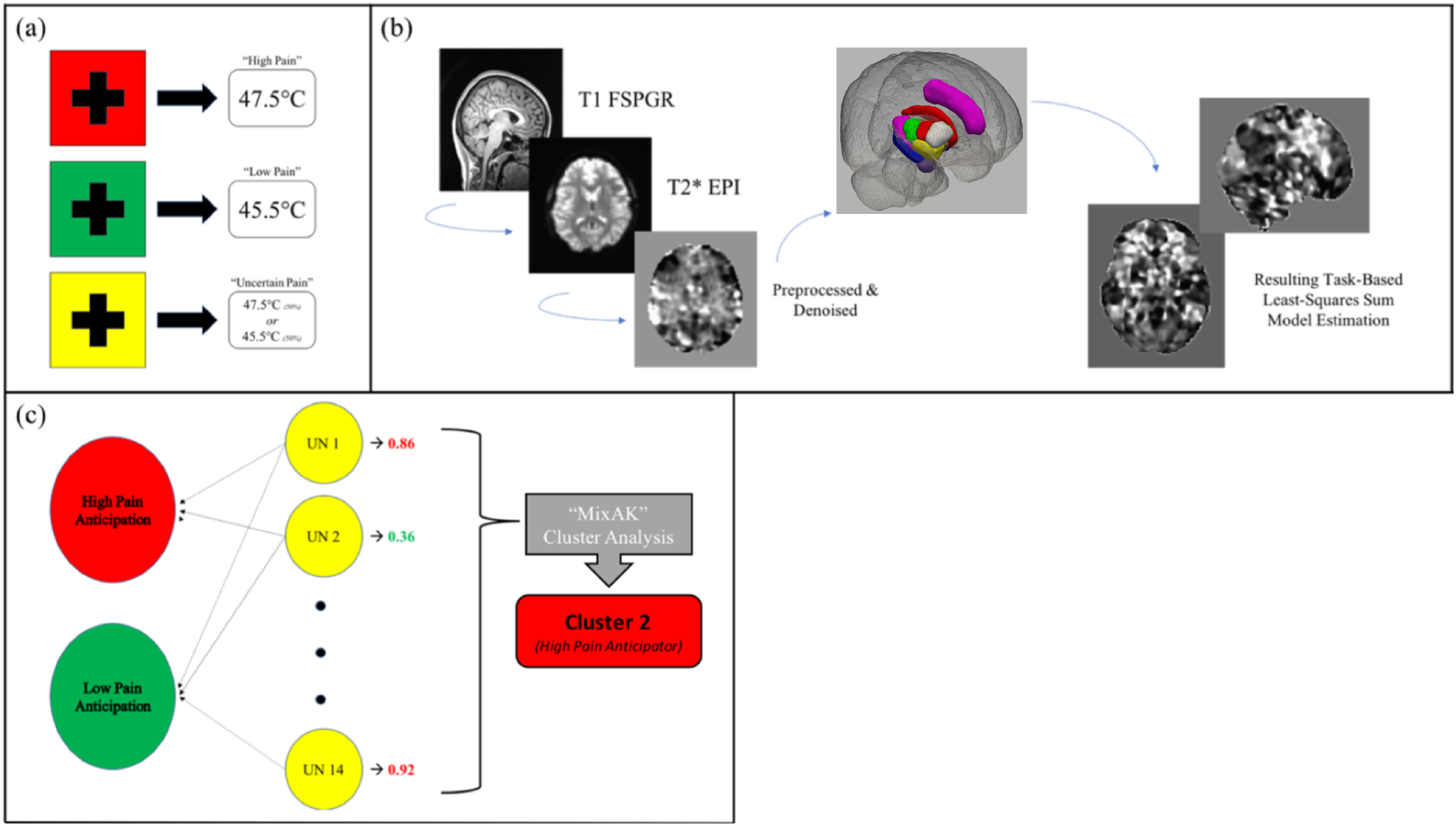
Illustration of applied methods. (a) Pain-anticipation paradigm; high pain (red, *N*=7), low pain (green, *N*=7), and uncertain pain (yellow, *N*=14) visual cues, followed by pain stimulation. (b) fMRI image pre-processing with CONN toolbox (www.nitrc.org/projects/conn, RRID:SCR_009550) and task-based regression (including least squares-sum model) completed in AFNI. Activation maps extracted in 26 *a priori* chosen ROIs depicted in glass brain on the right side only (c.f. Methods for more details) for each high and low pain anticipation event. (c) Each uncertain anticipation trial is compared to the certain activation maps and a probabilistic prediction is determined by LASSO. Predictions ≥0.5 are classified as “high”, and predictions <0.5 as “low”. Finally, predictions across all 14 uncertain trials for each subject are provided to mixAK cluster analysis in R, and each subject is clustered based on individual anticipation profile.

**Figure 2:**
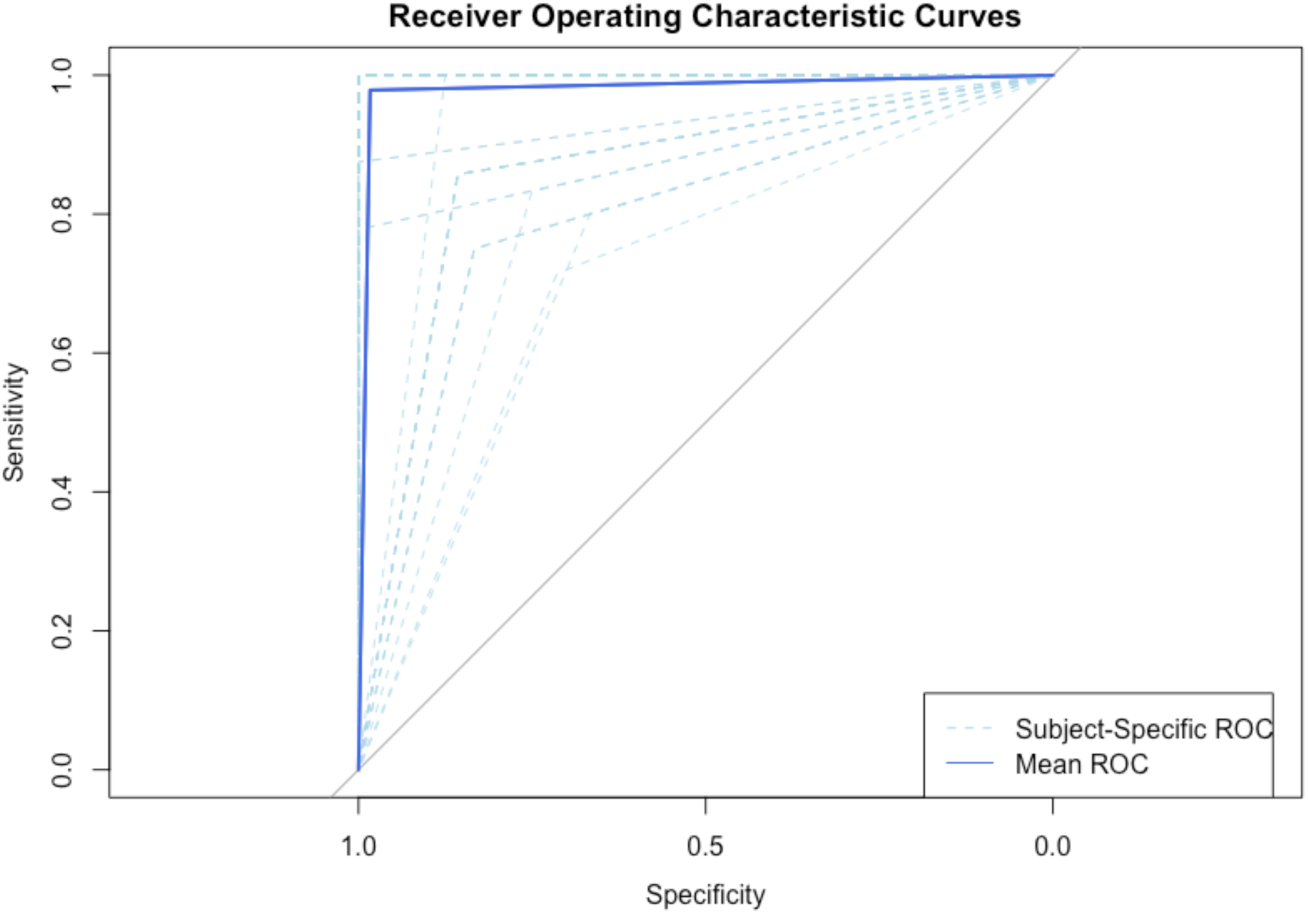
Receiver operating characteristic (ROC) curves of individual subject-specific LASSO models (dashed, light blue) in separating brain activity during anticipation of high-pain versus anticipation of low-pain and ROC curve of overall LASSO model performance within the entire study cohort (solid, blue).

To verify that the single-subject approach to distinguish between the high-pain and low-pain anticipatory neural patterns was the most effective method in this study, a population-based LASSO model was estimated based on population-wise activation maps averaged across all subjects and assessed to what extent population-based average pain-anticipatory neural patterns distinguished low-pain and high-pain anticipatory neural patterns on single-subject level. Population-based model performance was 59.4 ± 13.5% accurate across all subjects, with similarly low specificity, sensitivity, and AUC. Additionally, only twenty-two (15%) subjects’ high-pain and low-pain anticipatory neural patterns were separable at 75% accuracy level using the population-based LASSO model. This supports that population-based (or average) pain anticipation neural patterns are less reliable in distinguishing pain anticipatory neural patterns in every subject.

### Labeling each Uncertain Pain Anticipation Neural Response Pattern as Low-Pain or High-pain Anticipation in Every Subject

Once we confirmed that neural patterns of low pain and high pain anticipation are separable on individual level with high accuracy (see above and **Figure 2**), we then classified neural pattern during uncertainty into either high pain or low pain anticipation in each subject. The premise was that when presented with uncertainty, at each instance an individual would either anticipate the best-case scenario (i.e., positive bias, represented in this study by low-pain), or the worst-case scenario (i.e., negative bias, represented in this study by high-pain). Probed by each subject’s own individualized LASSO model for high-pain and low-pain anticipatory neural patterns separation, likelihood of low-pain versus high-pain anticipatory response was estimated for the subject’s brain activity patterns from each of the fourteen uncertain anticipatory trials, separately. Classifier decisions at each of the 14 anticipation periods during uncertainty were based on a continuous classifier evidence values (0-1) in that predictions ≥0.5 are classified as “high”, and predictions <0.5 as “low” (**Figure 3**).

**Figure 3:**
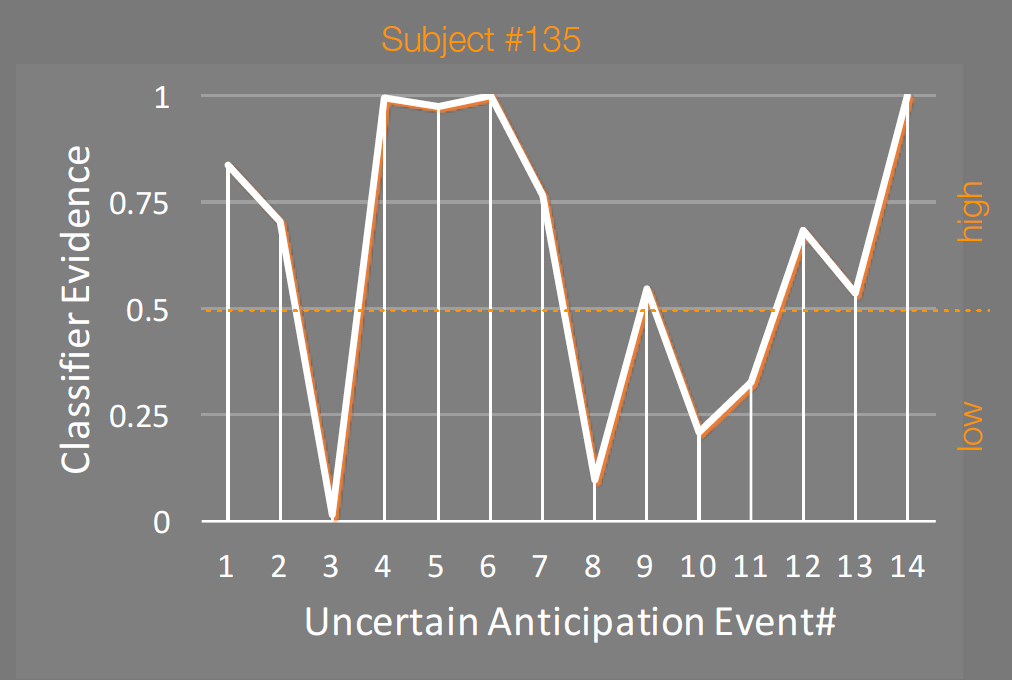
Uncertain anticipation decoding output for representative subject #135. Based on unique brain patterns learned from the certain pain anticipation conditions, each uncertain anticipation event is decoded by subject’s own individualized LASSO model. Output includes continuous classifier evidence values (0-1, y-axis) for each uncertain anticipation event (total of 14 events, x-axis). Classifier evidence indicates how brain pattern during each uncertain anticipation event matches the trained brain pattern for low (if <0.5) or high (if ≥0.5) pain anticipation pattern.

In order to ensure that we are classifying pain anticipation rather than the cue, we examined whether uncertain anticipation neural brain activity pattern differed from average certain anticipation neural brain activity pattern via a LASSO model. We found that certain (low-pain and high-pain) versus uncertain anticipation trials were only distinguishable with 45.8% accuracy. This finding suggested that BOLD response patterns in uncertain anticipation trials more closely resembled that of certain pain anticipation trials, as opposed to a separate pattern distinct to the uncertainty cue condition. In other words, the anticipatory pattern during uncertainty was related to the stimulus that followed the cue rather than the cue itself in isolation.

### Uncertain Pain Anticipation Pattern is Stable Across Time

To further validate our findings, stability of each subject’s decoded anticipation biases during uncertainty over time was assessed in a test-retest cohort of thirty-two subjects who had repeat fMRI data collected 12 ± 1 months apart. Each subject’s individualized LASSO modeling of certain anticipatory trials from the first imaging session (test) was applied to uncertain pain anticipation trials of the subject’s subsequent imaging session (retest). Twenty-one out of 32 subjects (65.6%) presented with the same decoded anticipatory biases (i.e., average likelihood of anticipating low-pain or high-pain during uncertain trials, quantitatively measured as average of subject’s prediction probabilities for uncertain trials across one imaging session) during both test and retest imaging sessions. Since test-retest cohort included individuals (n=24) who met criteria for major depressive disorder (MDD) at the initial fMRI session, we explored whether MDD diagnosis or current depressive symptom severity influenced uncertain anticipatory biases in these subjects. Eleven of the depressed subjects remitted by their final imaging session. We found that among the MDD subjects anticipatory biases were not influenced by their current diagnosis (t=0.13, p=0.90, df=57) or depressive symptom severity (t=0.98, p=0.33, df=57). These results point that pain anticipation response biases during uncertainty as determined by neural activity patterns are trait characteristics of an individual.

### Subjects Cluster by Anticipation Biases During Uncertainty

Our secondary aim was to discover whether anticipatory response biases during uncertainty may be influenced by psychopathology, as suggested by prior work using group-level analyses [21, 24, 25, 30–34]. We applied a generalized linear mixed model (GLMM) with Markov Chain Monte Carlo (MCMC) methods [35, 36] to perform an unsupervised clustering of subjects with similar decoded high-pain versus low-pain anticipatory response patterns to 14 uncertain trials. Effects accounted for in the GLMM included the fixed effect of trial time within the fMRI session and the random effect of the previous pain stimulation experienced (i.e., low-pain or high-pain). Previous pain was included as a random effect based on findings indicating that prior experiences can modulate the experience of painful stimulation [3, 4, 7–11].

Three clusters of subjects were identified based on subjects’ decoded response patterns to uncertain trials as shown in **Figure 4** (see Supplementary Material 5 for a cluster breakdown of demographic and neural activation factors). The first (“cluster 1”; blue) consisted of 55 subjects, all of whom had on average low-pain anticipatory response (i.e., positive bias) to uncertain trials. The second (“cluster 2”; red) had 47 subjects, all of whom had on average high-pain anticipatory response (i.e., negative bias) to uncertain trials. The remaining subjects (“unclassified”; gray) were unable to be classified into either of the two aforementioned clusters and all had an average low-versus-high likelihood of 0.25-0.75 across all uncertain trials, indicating that these subjects had highly variable anticipatory response patterns to uncertain trials over the course of the fMRI experiment. In accordance with this conclusion, there were statistically significant association between cluster and average prediction level (chi=98.8, *p*<0.01, df=2). Furthermore, as demonstrated in **Figure 5**, regardless of previous pain stimulation, cluster 1 subjects (blue) were significantly (chi=162.65, *p*<0.0001, df=1) more likely to anticipate low-pain (green), and cluster 2 subjects (red) were significantly more likely to anticipate high-pain (red). The unclassified subjects (grey) anticipated low- and high-pain at approximately 50% after high-pain, while after low-pain the unclassified subjects anticipated high-pain more often than low-pain.

**Figure 4:**
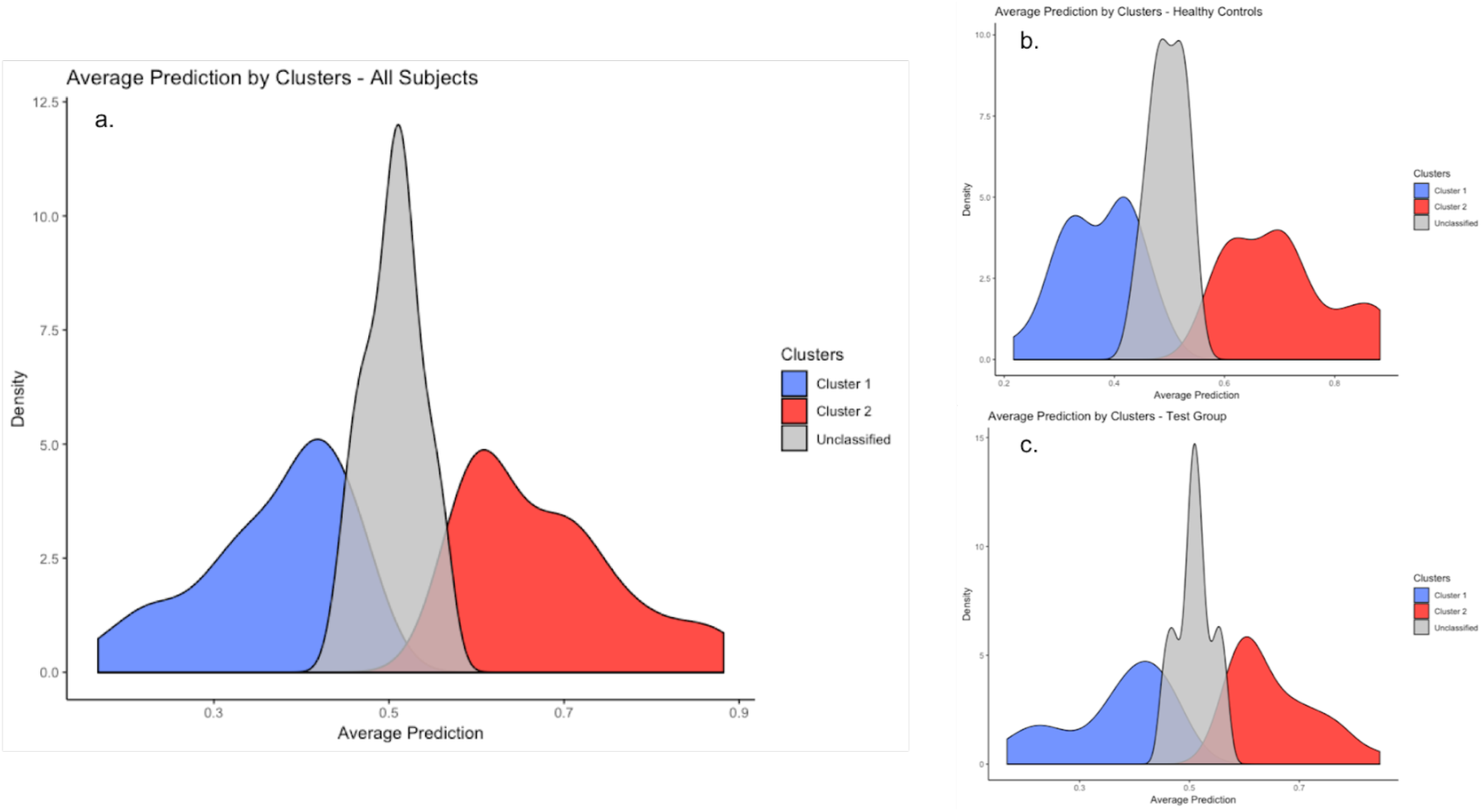
Histograms depicting cluster classification versus average predictions. Across all groups, (a) full cohort, (b) healthy controls, and (c) “mixed psychiatric” test group, subjects with average anticipation probabilities for uncertain trials less than 0.5 were classified in cluster 1, greater than 0.5 were classified in cluster 2, and close to 0.5 were unclassified. X-axis: average anticipation probabilistic prediction across 14 uncertain trials; y-axis: proportion of subjects in each cluster.

**Figure 5:**
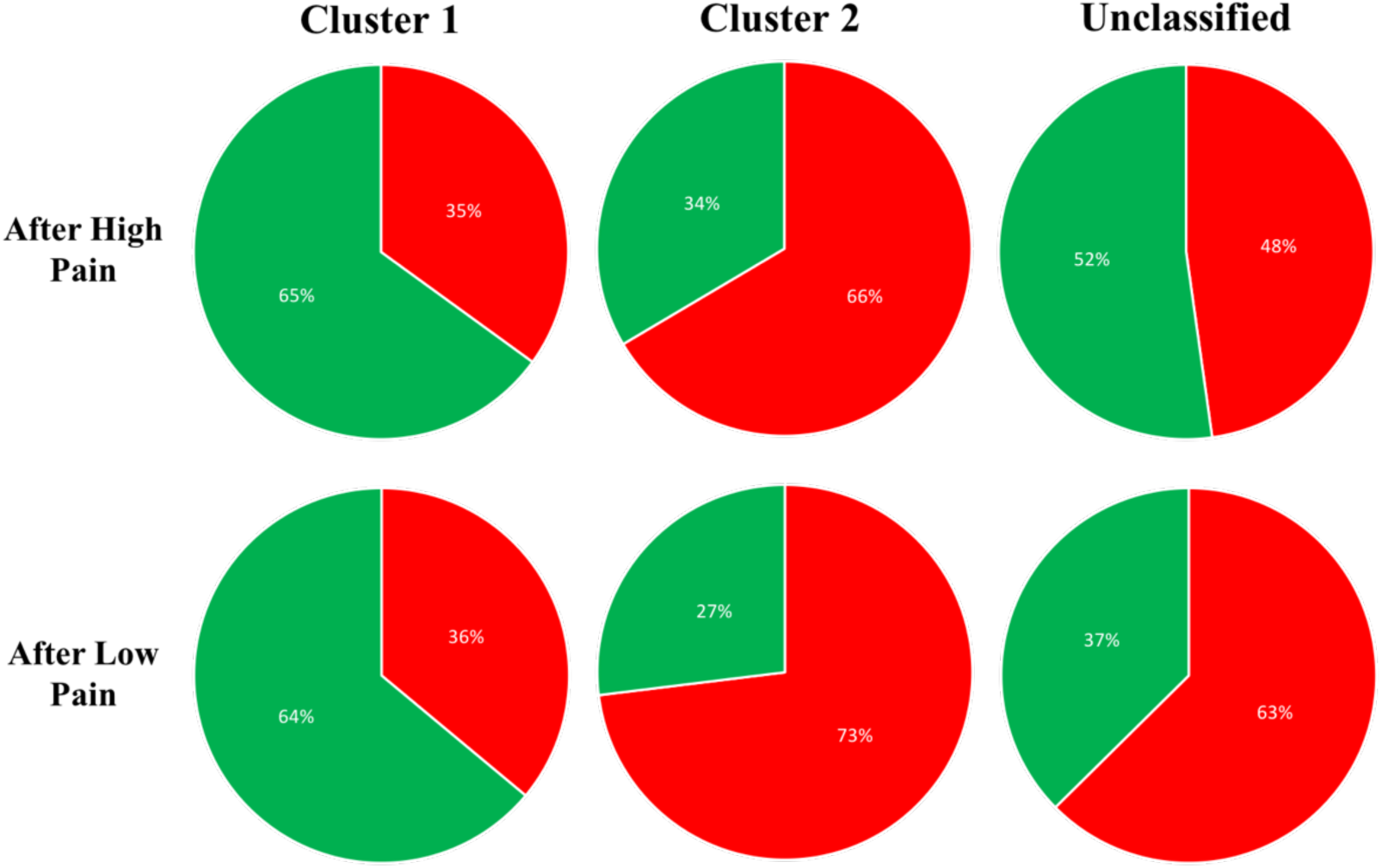
Uncertain pain anticipation predictions following low-pain and high-pain, separated by unsupervised cluster classification. Within reliable clusters the difference in predictions is not significant between two anticipation scenarios. Cluster 1 (low-pain anticipators, left) anticipate low-pain more frequently following both low- and high-pain stimulation. Cluster 2 (high-pain anticipators, middle) anticipate high-pain more frequently following both low- and high-pain stimulation.

**Figure 6:**
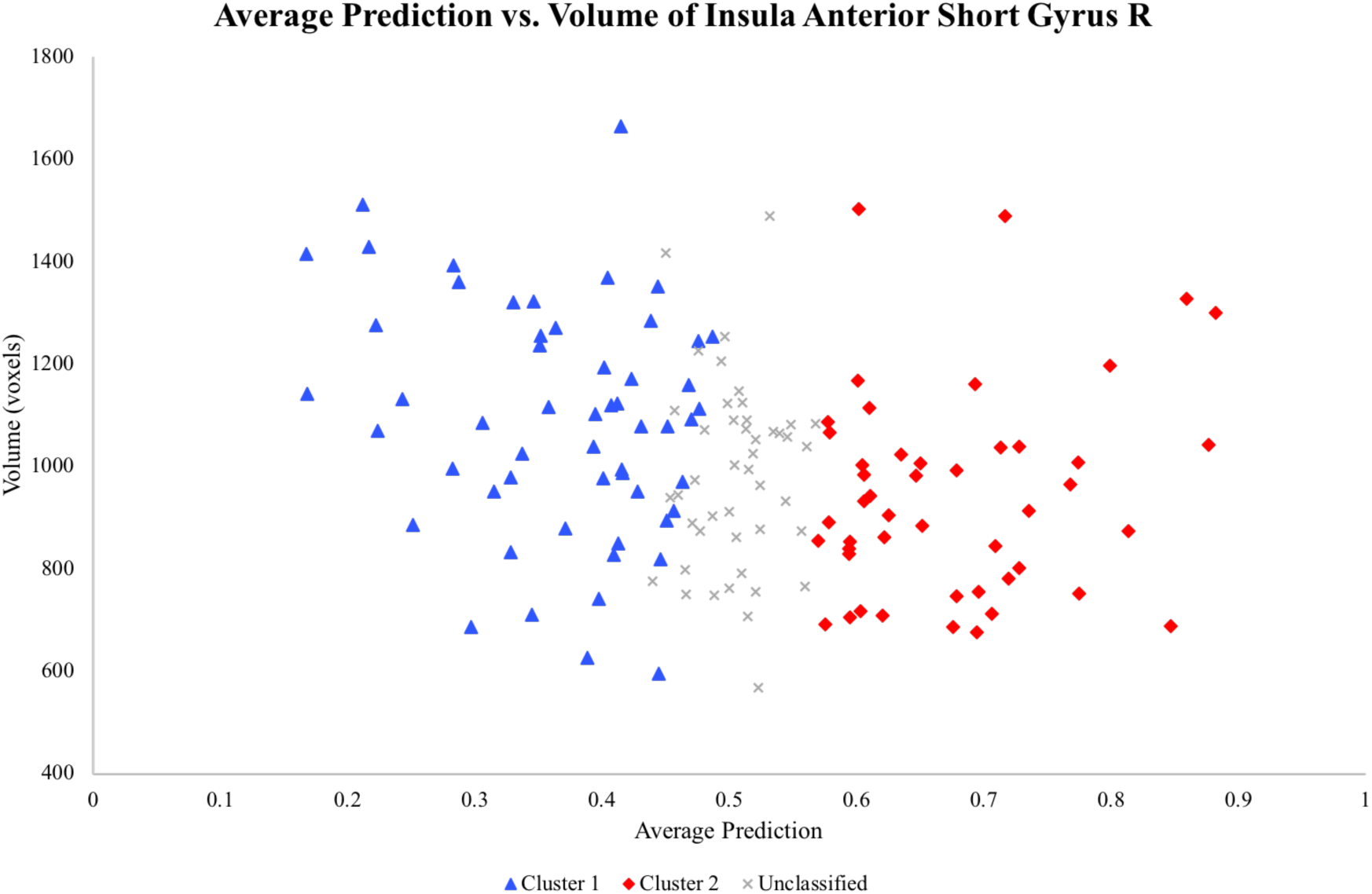
Scatterplot comparison of average prediction per subject versus the volume of the insular anterior short gyrus on the right hemisphere. Subjects are separated by unsupervised cluster classification. Symbols: Blue triangle, cluster 1; red diamond, cluster 2; gray cross, unclassified. x-axis: average anticipation probabilistic prediction across 14 uncertain trials; y-axis: volume of insular anterior short gyrus on the right hemisphere in voxels (1 × 0.97 × 0.97 mm^3^). (Abbreviations: R, right cortical hemisphere.)

Contrary to our hypothesis, subjects did not reliably cluster based on group (healthy vs. “mixed psychiatric”), clinical diagnosis (see Supplementary Material 7 for non-significant trends), age, or gender. Of note, due to the nature of the “mixed psychiatric” cohort, these individuals had many overlapping psychiatric symptoms rather than representing a “clean” DSM-IV diagnosis. It may thus be possible to separate subjects based on their anticipatory bias during uncertain trials in a larger number of subjects with a single psychiatric symptom. Nevertheless, we did not find any significant relationship between current depressive symptoms (t=0.51, p=0.61, df = 145; measured by BDI-2) or pain cognitions (t=0.37, p=0.71, df = 125; measured by pain catastrophizing scale, PCS; *N*=20 subjects (8 healthy controls and 12 “mixed psychiatric” did not complete PCS scale) and uncertain pain anticipation predictions.

### Anticipatory Response Biases during Uncertainty are Related to Brain Structure

To assess to what extent the anatomic architecture was associated with the decoded anticipatory response biases during uncertainty, we first estimated gray matter tissue volume of each of the regions of interest for each subject using Advanced Normalization Tools (ANTs) [37]. To control for the total intracranial volume, each regional volume estimates was normalized per subject by the determinant of the affine transformation (i.e., the relative amount of global shape scaling required to transform a subject’s brain to the MNI template space). Out of twenty-six regions-of-interest assessed in the study, only the volume of the anterior short insular gyrus on the right hemisphere was significantly and inversely associated with the average decoded anticipatory response bias during uncertain trials (**Figure 7**, Pearson correlation coefficient r=-0.33, *p*=0.001, df=145), suggesting that those with the greatest right anterior short insular gyrus volume were more likely to anticipate low-pain during uncertainty (i.e., more likely to show “positive bias” and be classified within cluster 1). Consistent with this, the right anterior short insular gyrus volume significantly differed across three clusters (ANOVA, *F*=6.72, *p*=0.002, df=146).

Although not significantly correlated to average probabilistic decoding of the anticipatory response during uncertain trials, the anterior short insular gyrus on the left hemisphere (Pearson correlation coefficient r=-0.302, *p*=0.003, df=145), and the anterior cingulate cortex on the left hemisphere (Pearson correlation coefficient r=-0.31, *p*=0.002, df=145), showed strong trends compared to anticipation biases during uncertainty.

## Discussion

In this study, we employed innovative single-subject machine learning techniques to fMRI data from an evoked pain anticipation paradigm to study individual differences in pain anticipation behaviors. Our major findings are as follows: 1) Using a multivariate pattern analysis and a single-subject analytics it is possible to distinguish between neural activation patterns of the anticipation of high and low pain stimulation on single subject level with high accuracy, sensitivity and specificity. This is novel and to the best of our knowledge has not been demonstrated before. 2) There is high variability in functional neural activation patterns among subjects that contributed to pain anticipation classification emphasizing the need for single-subject approach. 3) Based on a data-driven unsupervised clustering approach, individuals have intrinsic neural anticipatory patterns distinctly separating them as adaptive (i.e., positive bias) or maladaptive (i.e., negative bias) pain anticipators, a behavior that does not reliably map onto mental health status or other common demographics, but rather is more influenced by the underlying anatomical architecture. These findings add valuable information to the current models of pain processing and predictions. Our findings support that anticipation behavior is specific to each individual, stable over time, and potentially less dependent of subjective psychiatric diagnosis.

This is the first study to demonstrate that there is a distinguishable subject level difference between neural activation patterns of differing levels of pain anticipation. Previous studies have used machine learning to discriminate between (1) painful heat and non-painful warmth stimuli based on subject ratings post-stimulus, and (2) pain anticipation and pain recall [28], using group-wise analyses as opposed to the single-subject approach used here. Additionally, we show that activation patterns during uncertain anticipation more closely resemble that of certain possible outcomes, as opposed to a distinct uncertainty pattern. This asserts that following uncertain or ambiguous cues individuals rely on prior knowledge to predict the possible outcome and could in turn demonstrate a bias toward preparing for the worst case or hoping for the best case (i.e., expecting pain or looking forward to relief). This bias was also shown to be consistent within most individuals irrespective of the previous pain experience, which indicates stability within the experimental paradigm setup. Using multi-variate pattern analysis (MVPA) and LASSO machine learning techniques, these differences were seen at the level of the insula, anterior cingulate, amygdalae, and striatum. The regions-of-interest (ROIs) chosen for this study were previously implicated in multidimensional pain experience [1, 19, 21, 25, 28–30] but we found that the predictive value of each region for low-versus-high pain anticipation varied widely across subjects. The most common variables predictive of low-versus-high anticipatory responses included the insular regions, particularly the anterior short gyrus (functionally and structurally), and the nucleus accumbens (functionally only) with varying inclusion frequencies in the individualized LASSO models. This emphasizes the immense biological heterogeneity present within the study cohort, and ultimately the importance of single-subject analytic approaches to capture this variability.

The overlap between functional and structural underpinning of anticipatory response supports previous suggestions to integrate structural and functional data to fully understand neurological processes [38]. In doing so we can better develop a full picture of how a cortical region, like the insular anterior short gyrus, might be crucial to pain anticipation and processing [19]. Additionally, the frequent inclusion of the nucleus accumbens, which has previously been implicated in the anticipation of pain relief [39] and in animal studies of pain-predictive cue-influenced decision making [12], suggests that there is a synergistic relationship between the anticipation of pain and the expectation, or hope, of relief from such pain. We posit that in comparison to the high-pain, subjects feel a sense of pain relief when experiencing the low-pain which can motivate anticipation. Our finding that activation during uncertain-versus certain-anticipations were not distinguishable indicates that humans will inherently make predictions about future events given their past experience or understanding of possible outcomes. It appears that the underlying mechanism of pain anticipation may actually be a tradeoff between the anticipation of a negative stimulus versus a positive outcome, which supports the motivation-decision model of pain proposed by Fields [13].

We were interested in testing whether subjects’ predictions in the face of uncertainty would cluster according to their DSM-IV diagnoses or would be influenced by demographic variables such as gender and age, all of which have been reported to associate with response to uncertainty [21, 24, 25, 30–34] using group level analyses. Using unsupervised clustering methods, we found that our study subjects were grouped solely based on their anticipation bias patterns over the course of the entire pain-anticipation paradigm, rather than the presence of a mental illness as it has been defined by the DSM-IV. As for those subjects who were successfully clustered, cluster 1 represented the low-pain anticipation group, and cluster 2 the high-pain anticipation group. Most compelling were the differences between clusters in anticipatory behaviors following low- and high-pain stimulation. The two distinguishable clusters (cluster 1 and 2) were consistent in their anticipation of low- and high-pain, respectively, regardless of the previous stimulus level. The same was not true for the remaining subjects who could not be reliably classified into the low- or high-pain anticipator clusters. These subjects showed equal anticipation of low- and high-pain following high-pain stimulation, yet a greater instance of high-pain anticipation following low-pain. These findings might suggest that there are individuals who are consistent in their anticipation patterns during uncertainty, and there are subjects whose anticipation is more clearly influenced by prior experiences. Since subjects were not clustered by group, gender, or DSM-IV diagnoses, further work should be completed to understand what differentiates these influenceable subjects from those whose anticipatory responses are more rigid.

While there was no significant difference in unsupervised cluster classification between the healthy controls and “mixed psychiatric” test group, analysis of the diagnostic subgroups within the test group had interesting observations that merit further study (see Supplementary Material 7). For instance, although limited by small sample sizes, it is still noticeable that within recovered anorexic females and combat-exposed males suffering from post-traumatic stress disorder (PTSD) and mild traumatic brain injury (mTBI), there is a propensity for subjects to anticipate low-pain or high-pain, respectively (**Supplementary Figure 3**). The criteria for these disorders, including low body mass index and a significant traumatic experience as result of brain injury, are both quantifiable and thus potentially limit the amount of heterogeneity within diagnostic groups. The same cannot be said for subjects diagnosed with MDD, whose diagnosis relies more on subjective measures, and thus highlights a pressing concern of high heterogeneity within the diagnostic label which necessitates methods for identifying these differences to more effectively treat all individuals. For MDD, and many other psychiatric disorders, common treatment calls for the use of cognitive behavioral therapy (CBT). The main aim of which is to identify, challenge, and change dysfunctional cognitions and behaviors, and to equip patients with effective coping strategies. Therefore, understanding how an individual perceives the world objectively can allow for CBT treatments to be tailored on a patient-by-patient basis. Objective measures also remove limitations of patient self-report that are often limited in vulnerable populations (such as those with mental illness). Additionally, it has been shown that subjective self-report of pain is strongly influenced by relative experiences, such that a moderate pain stimulus may be rated as painful when paired with a neutral, non-painful, stimulus, but when contrasted with an intense pain stimulus the moderate pain may be rated as “pleasant” [3]. Using a pain-anticipation paradigm like the one employed in this study would be especially useful as it avoids issues of self-report, instead capturing implicit behavioral responses.

After determining that we could classify subjects based on their anticipation biases during uncertainty, we wanted to know if these individualized classifications were stable over time. Findings from the test-retest group confirm our conclusion that anticipation is an innate trait as opposed to a state-based phenomenon. If stability of anticipation biases had only been seen in subjects whose mental health remained stable (healthy controls or MDD without entering remission or relapsing) then it would have indicated an important effect of current mental state on anticipation behaviors. Instead, the majority of subjects, regardless of initial diagnosis or current mental health state, remained stable in their average prediction over the course of two or three imaging sessions. It is recommended that future studies conduct follow-up imaging sessions with a larger cohort comprised on a greater variety of diagnoses.

One limitation to this study is the difference in the proportion of males and females overall. The role that gender plays in anticipation and the perception of pain has been investigated previously, and it has been found that females tend to experience higher levels of psychological distress than males in response to painful stimulation [34, 40]. Although our results did not clearly separate anticipation patterns between genders, we found that females in the “mixed psychiatric” test group on average tended to anticipate high-pain more frequently than healthy females (*p*=0.099). We believe that a larger cohort, with an equal number of participants from each gender, might be instrumental to better assess potential gender-differentiated response patterns in which females, especially those with current or past psychopathology, anticipate high-pain more frequently than males.

## Conclusion

The distinct anticipation bias clusters identified here represent the intrinsic proclivities specific to each individual to either see the world through an optimistic or pessimistic filter. With state-of-the-art machine learning techniques, this study demonstrated with high accuracy the ability to distinguish between neural activation patterns of low- and high-pain anticipation on a single-subject basis, and label uncertain anticipation neural activity accordingly. This ability was formed on the finding that when presented with an uncertain pain cue, humans inherently anticipate based on known possible outcomes. Further, it was shown that an individual’s neurobiological anticipation signature is unique to them, most often stable over time, and relates more strongly to the underlying anatomy than clinical diagnosis. This variation in anticipation behaviors and regional activation maps emphasizes the need for single-subject based machine learning analyses in order to avoid over-generalizations. It is our hope that techniques such as those presented in this study will eventually be applicable in clinical settings to ascertain objective measurements of intrinsic behaviors.

## Methods

### Participants

Data for one hundred and forty-seven subjects (50 females, mean ± SD age, 28 ± 6.8 years) was used for the current study. Subjects were recruited using flyers at University of California San Diego (UCSD) clinics, internet sites (e.g., Craigslist), local papers, and word of mouth. The study was approved by the UCSD Human Research Protection Program and Veterans Affairs San Diego Healthcare System Research and Development Committee. Prior to participating all subjects gave their written informed content and underwent a Structured Clinical Interview administered by trained interviewers according to the Diagnostic and Statistical Manual for Mental Disorders (DSM)-IV [41] to establish current and past psychiatric diagnoses. Fifty-seven subjects (22 females) were healthy controls with no current or past history of mental illness or trauma, and ninety (28 females) were part of the test group (“mixed psychiatric”). Subjects completed behavioral questionnaires, including the Beck Depression Inventory (BDI)-2 [42] for depressive symptom severity. Subjects were excluded from the study if they: (1) used psychotropic medication within the last 30 days; (2) fulfilled DSM-IV criteria for alcohol/substance abuse or dependence within 30 days of study participation; (3) fulfilled DSM-IV criteria for lifetime bipolar or psychotic disorder; (4) has ever experienced a head injury; (5) had clinically significant comorbid medical conditions, such as cardiovascular and/or neurological abnormality, or any active serious medical problems requiring interventions or treatment; (6) had a history or current chronic pain disorder; (7) had irremovable ferromagnetic material; (8) were pregnant or claustrophobic; and (9) were left-handed. All female subjects were scanned during the first ten days of their menstrual cycle. A subset of these subjects (N=32) underwent the same pain-anticipation paradigm on at least two separate occasions. Each scan was roughly 12 months (± 1 month) apart. Eleven completed three scanning sessions. This cohort will be referred to as the “test-retest” subjects.

### Neuroimaging Protocol

Two fMRI runs (412 brain volumes per run) sensitive to BOLD contrast were collected for each subject using a 3.0 Tesla GE Signa EXCITE scanner (GE Healthcare, Milwaukee, WI, USA) (T2*-weighted echo planar imaging, TR=1500ms, TE=30ms, flip angle=90, FOV=23cm, 64 × 64 matrix, 30 2.6-mm 1.4-mm gap axial slices) while they performed the pain-anticipation paradigm described in Supplementary Material 1. Acquisitions were time-locked to the onset of the task. During the same experimental run, a high-resolution T1-weighted image (FSPGR, TR=8ms, TE=3ms, TI=450ms, flip angle=12, FOV=25cm, 172 sagittal slices, 256 × 256 matrix, 1 × 0.97 × 0.97 mm^3^ voxels) was obtained for anatomical reference. The fMRI protocol was the same for the majority (*N*=125) of the subjects, but a small subset of subjects (11 female recovered anorexics and 11 female healthy controls) were scanned separately with a TR of 2000ms for the T2*-weighted echo planar imaging. The change in timing was controlled for during preprocessing so that all following analyses were completed in unison.

### Pain-Anticipation fMRI Paradigm

The pain-anticipation paradigm used two predetermined and consistent temperatures, 45.5°C and 47.5°C, across subjects to elicit mild (“low-pain, LP”) and moderate (“high-pain, HP”) sensations, respectively. Stimulation was delivered through a 9cm^2^ thermode (Medoc TSA-II, Ramat-Yishai, Israel) on the participant’s left forearm, as described elsewhere [24]. Each trial began with a period of anticipation initiated by a visual cue (**Figure 1a**). The cue was always followed by painful stimulation (either HP or LP), and a period of rest (jittered between 24-30s) before the next trial began. The schedule of stimuli differed between runs in a pseudorandom and counterbalanced order (see Supplementary Material 1). A single imaging session include 7 HP trials (HP cue followed by HP stimulation), 7 LP trials (LP cue followed by LP stimulation), and 14 uncertain (UN) trials (nonspecific pain cue followed by either HP or LP stimulation at 50% probability). For a more detailed explanation of the pain-anticipation paradigm, see Supplementary Material 1.

### fMRI Image Processing

All fMRI data was preprocessed using a MatLab-based functional connectivity toolbox, CONN [43], to denoise and align the images for analysis. A detailed account of the preprocessing pipeline is given in Supplementary Material 2. Further analysis was conducted using the Analysis of Functional NeuroImages (AFNI) software package [44]. A multiple regression model corrected for autocorrelation consisting of twenty-eight anticipation-related regressors and twenty-eight stimulus-related regressors was applied to preprocessed time-series data for each individual. A separate regressor was calculated for each trial such that each event had its own estimated amplitude. The 28 anticipation-related regressors modeling the entire anticipation period consisted of: (1) anticipation of moderately painful heat stimulation (HP anticipation, *N*=7), (2) anticipation of mildly painful heat stimulation (LP anticipation, *N*=7), and (3) anticipation of uncertain painful heat stimulation (UN anticipation, *N*=14). All stimulation conditions (14 HP and 14 LP) were modeled as regressors of no interest. Six additional regressors were included in the model as nuisance regressors: one outlier regressor to account for physiological and scanner noise (that is, the ratio of brain voxels outside of 2 standard deviations of the mean at each acquisition), three movement regressors to account for residual motion (in the roll, pitch, and yaw, directions) and regressors for baseline and linear trends to account for signal drifts. To reduce the false positives induced by cross-correlations, time-series data were fit using the AFNI [44] program 3dLSS [45]. 3dLSS applies a least-squares-sum model estimation to the resulting individually modulated time series data in order to deconvolve BOLD activation in the MVPA of task-based fMRI data [45]. This approach was chosen following previous findings that 3dLSS, as opposed to a general linear model (GLM) achieved higher classification accuracy with low variance [45].

### Structural Analysis

Advanced Normalization Tools (ANTs) [37] were used to extract geometric measurements for each ROI per subject. Only the volume of each ROI was of interest in the current study. To extract volumes (measured in voxels), a combination of ANTs scripts (“antsRegistrationSynN” and “antsApplyTransforms”) [37, 46] were used. For volumetric analysis, the volume of each region was normalized per subject using the determinant of the affine transformation (i.e., the relative amount of distortion required to transform a subject’s brain to the MNI template space).

### Regional Activation Maps

Activation maps were created on a single-subject basis. Masks of selected ROIs were created in MNI space using AFNI and were applied to the functional activation maps using 3dDeconvolve [26]. A total of twenty-six ROIs were chosen based on their prominent roles in pain prediction, processing and relief [13, 26, 27]. Twelve ROIs (6 on each side) were selected within the insula: (1) posterior long gyrus, (2) anterior short gyrus, (3) middle short gyrus, (4) posterior short gyrus, (5) anterior inferior cortex, and (6) anterior long gyrus (**Figure 1b**). Additionally, fourteen functionally relevant bilateral ROIs (7 on each side) were selected: (1) anterior cingulate cortex, (2) amygdala, (3), nucleus accumbens, (4) caudate nucleus, (5) putamen, (6) pallidum, and (7) substantia nigra. The anterior cingulate and amygdala ROIs were chosen for their role in affective processing networks, the nucleus accumbens as representative of the ventral striatum, along with its common targets, the pallidum and substantia nigra, and lastly the caudate nucleus and putamen as representative of the dorsal striatum. Using a separate AFNI program, 3dROIstats [44], the mean activation was extracted as the beta coefficient from each region during each anticipation trial. The use of t-values, as opposed to beta-coefficients, has improved classification accuracy in previous studies [47]. T-statistics were thus calculated in each region such that the t-statistic is equal to the beta coefficient divided by the standard error [28].

### Functional Analysis of Regional Activation Maps

Average activation within each region underwent regression analysis by way of LASSO. The LASSO regression model was executed in R [48] using the glmnet package [48] for Lasso and Elastic-Net Regularized General Linear Models. LASSO was performed on a single-subject basis in which the training set was individuals’ neural activation (from 26 ROIs simultaneously as independent predictors) during 14 certain anticipation trials (HP and LP as dependent outcome), and the test set was neural activation during the 14 uncertain trials. Logistic regression methods such as LASSO are especially important in this case as they allow for a smaller number of predictors to be included in the model. In glmnet, two variables, alpha and lambda, must be specified to control the fit and regularization of the LASSO model. Alpha represents the elastic-net mixing parameter such that a value of 0 uses a ridge penalty, 1 uses a LASSO penalty, and an intermediate value uses a weighted combination of the two. Lambda is the regularization parameter. Per subject the values for alpha and lambda were optimized prior to regression. The optimal value of alpha for each subject was determined by testing values (0-1) at regular intervals of 0.1 and selecting that which resulted in the greatest subject-specific classification accuracy. Regression was then fit to the training set and, given the optimal alpha, lambda was optimized by cross-validating the model one hundred times, averaging the error curve, and selecting a lambda associated with the minimum of the error curve. This allowed for accurate and consistent discrimination between single-subject neurobiological patterns of low- and high-pain anticipation.

LASSO predictions were made on the test set at a probabilistic level, based on the correlation of the activation t statistic to the cross-validated glmnet model. The predicted anticipation of each of the fourteen uncertain trials was calculated based on regional activation and recorded separately for each participant.

### Cluster Analysis

Cluster analysis was completed using the R-based package MixAK [35, 36]. Using MixAK we applied a generalized linear mixed model (GLMM) with Markov Chain Monte Carlo (MCMC) methods [35, 36] to cluster subjects based on the multidimensional (i.e., N=14 uncertain trials separately) probabilistic predictions made for each UN trial obtained from the LASSO model. Effects accounted for in the GLMM included the fixed effect of time and the random effect of the previous pain stimulation experienced (HP or LP). Previous pain was included as a random effect based on findings indicating that prior experiences can modulate the experience of painful stimulation [3, 4, 7–11]. The model used a normal mixture distribution of random effects. The maximum number of mixture components was set to four.

### Functional Analysis of Test-Retest Subjects

For the subset (*N*=32) of the cohort that underwent multiple scanning sessions, LASSO was completed with data from the follow-up sessions to assess the stability of the model and of prediction profiles over time. Setup for the LASSO model was consistent across all sessions per subject. The model used for prediction analysis was trained on the expected anticipation activation maps from session 1 for each subject and applied to all subsequent sessions.

### Statistical Analysis

To explore whether psychiatric and demographic variables influenced predictions, we performed chi-square tests on the LASSO prediction analysis results to compare predictions between the healthy controls and mixed psychiatric group, as well as males and females. Pearson correlation tests were performed to assess possible relationships between cluster classification and demographic variables (sex and age), as well as psychological variables (BDI, PCS, and comorbid diagnoses). For cluster analysis, chi-square tests, two-tailed t-tests, and an ANOVA, were also run.

## Author Contributions

I.S., D.T., and M.K., conceived and designed the analysis; I.S. collected the fMRI and behavioral data; D.T. and I.S. contributed tools for analysis; M.K. performed the processing and analyzed the data; I.S., D.T., and M.K. wrote the paper.

## Competing Interests

The authors declare no competing interests.

