## Supplemental Material for "BOLD Decoding of Individual Pain Anticipation Biases During Uncertainty"

### Supplementary Figures

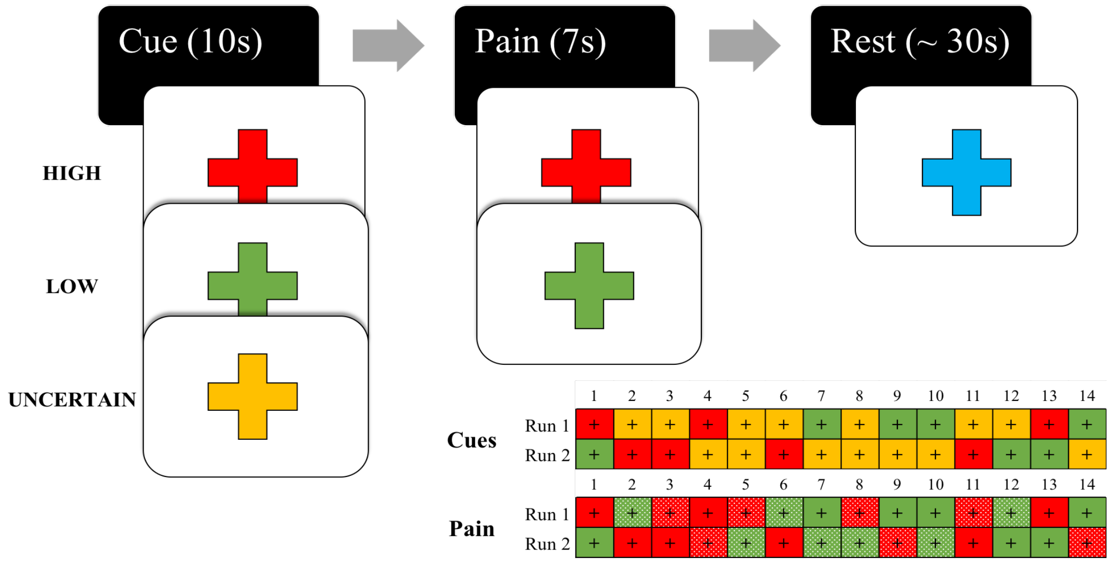


**Supplementary Figure 1: Schedule of pain-anticipation paradigm.** Simple breakdown of ordered pain-anticipation-rest timeline. Cues appear as presented to subjects. RED represents high-pain cue followed by high-pain administration (HP). GREEN represents low-pain cue followed by low-pain administration (LP). YELLOW represents uncertain pain cue followed by either high or low-pain administration (UN). Bottom right: timeline for one complete imaging session (both run 1 and run 2) separated by anticipation cue timeline (HP, LP, and UN), and pain stimulation (HP and LP).

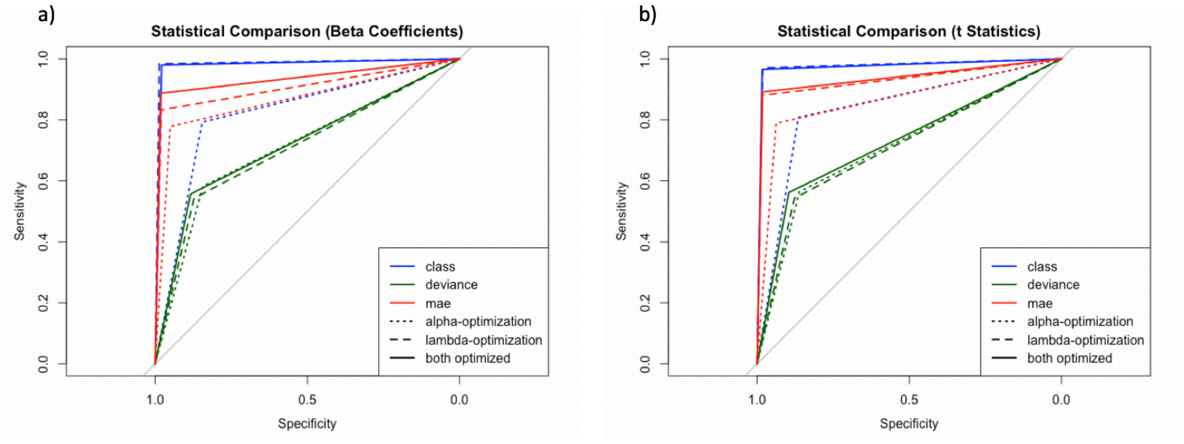


**Supplementary Figure 2: Receiver Operating Characteristic (ROC) curves for LASSO model setup results.** (a) beta-coefficient activation maps and (b) t-statistic activation maps. Loss parameter used in the cross-validation step was varied between trials such that; "class": misclassification error; "deviance": squared error; "mae": mean absolute error.

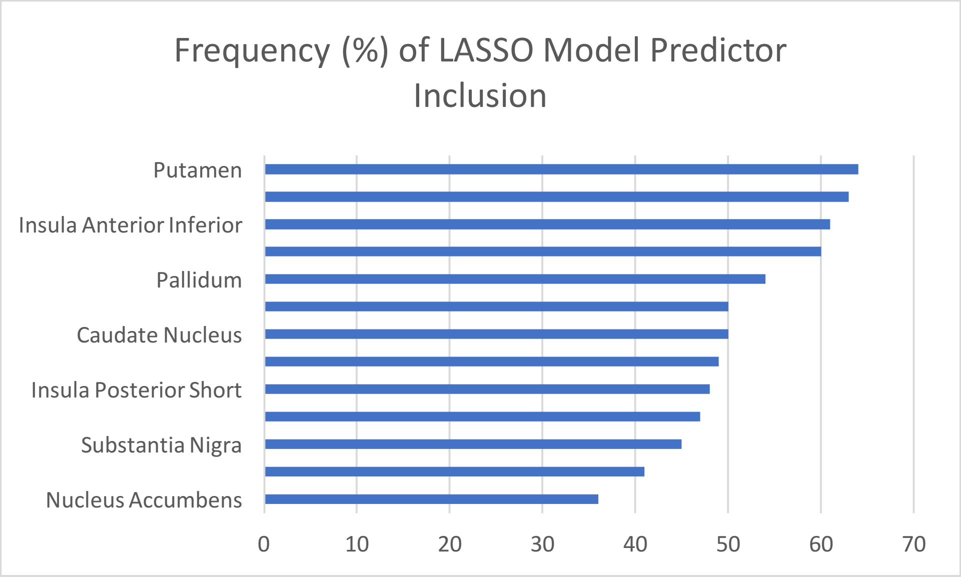


**Supplementary Figure 3: Frequency map of regions of interest included in single-subject LASSO models.** Models varied between subjects, resulting in a high variation of predictors included in each model. X-axis is frequency (in percent) of inclusion in all single-subject LASSO models (N=147). (Abbreviations: LASSO, least absolute shrinkage and selection operator)

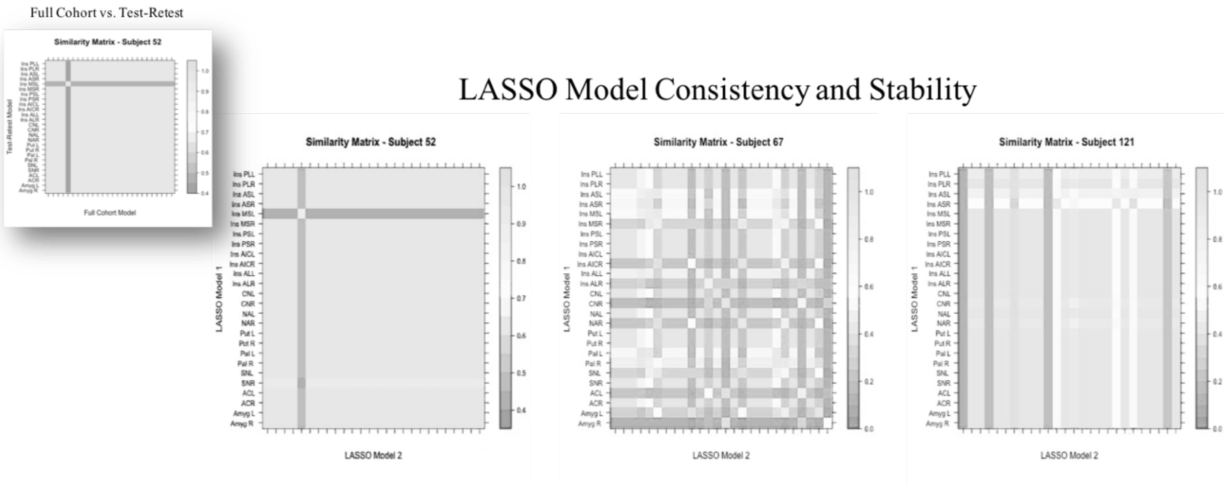


**Supplementary Figure 4: Similarity matrices comparing two runs (test-retest) of the LASSO model for three sample subjects.** The LASSO model from run 1 is represented on the vertical axis and the model from run 2 is on the horizontal axis. White indicates high similarity (equal, or close to equal, weight in the two LASSO models) and gray indicate large differences. Inset: Subject 52 (left) similarity matrix for LASSO model with full cohort (x-axis) and test-retest cohort (y-axis). (Abbreviations: LASSO, least absolute shrinkage and selection operator.)

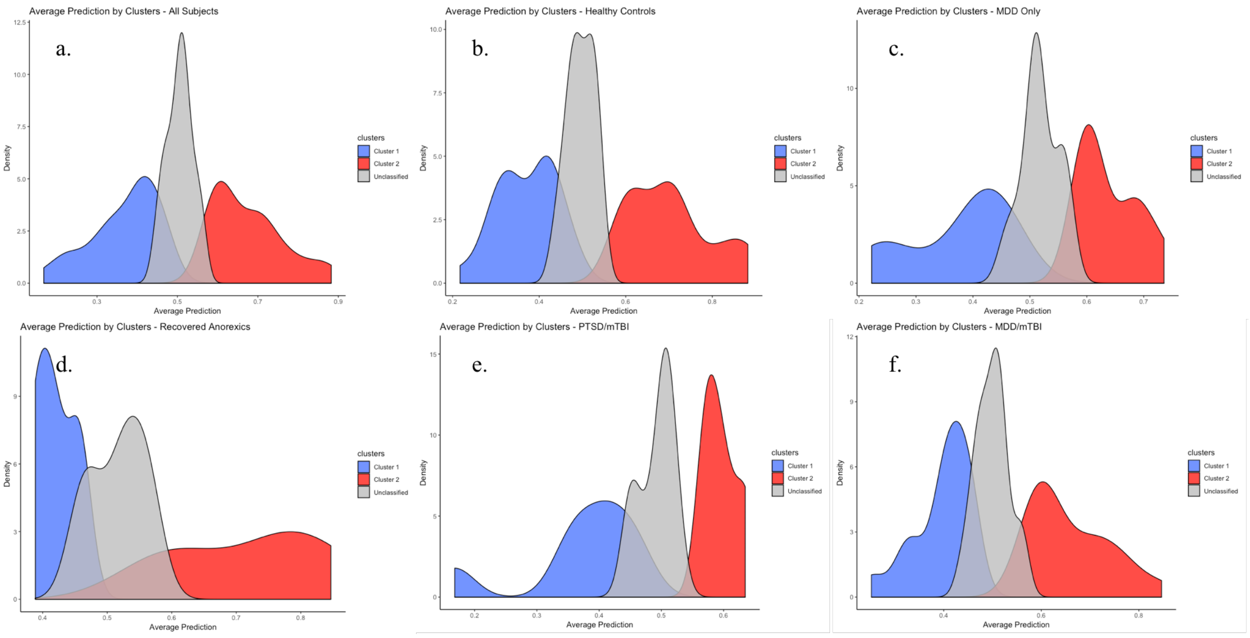


**Supplementary Figure 5: Cluster classification breakdown within groups.** Groups: (a) full cohort, (b) healthy controls, (c) MDD-only, (d) recovered anorexia, (e) PTSD/mTBI, (f) MDD/mTBI. (Abbreviations: MDD, major depressive disorder; PTSD, post-traumatic stress disorder; mTBI, major traumatic brain injury.)

### Supplementary Tables

| | 57 Healthy Controls | 90 Mixed Psychiatric | Stats | |
|---|---|---|---|---|
| | *Mean* | *Mean* | *t* / $\chi^2$ | *p* |
| **Demographic Variables** | | | | |
| *Gender* | 35M/22F | 62M/28F | 0.87 | 0.35 |
| *Age (years)* | 27.3 ± 7.4 | 28.4 ± 6.4 | -0.97 | 0.33 |
| *Race* | | | 0.64 | 0.88 |
| *African American* | *N*=4 | *N*=5 | | |
| *Asian* | *N*=7 | *N*=8 | | |
| *Caucasian* | *N*=30 | *N*=49 | | |
| *Other* | *N*=16 | *N*=28 | | |
| **Clinical Variables** | | | | |
| *Major Depressive Disorder* | | *N*=31 | | |
| *Combat Trauma* | | *N*=19 | | |
| *mTBI* | | *N*=44 | | |
| *Recovered Anorexia* | | *N*=11 | | |
| **Psychological Variables** | | | | |
| *Beck Depression Inventory-2* | 1.9 ± 3.1 | 17.9 ± 12 | -9.75 | **<0.001** |

**Supplementary Table 1: Demographic, clinical, and psychological variables of the full cohort.** (Abbreviations: F, female; M, male; mTBI, mild traumatic brain injury.)

| | 8 Non-MDD | 24 MDD | | |
|---|---|---|---|---|
| | *Mean* | *Mean* | $t / \chi^2$ | *p* |
| **Demographic Variables** | | | | |
| *Gender* | 8M/0F | 17M/7F | 1.16 | 0.28 |
| *Age (years)* | 27.1 ± 4.14 | 26.9 ± 7.2 | -0.07 | 0.94 |
| *Race* | | | 1.08 | 0.78 |
| *African American* | *N*=4 | *N*=13 | | |
| *Asian* | *N*=2 | *N*=9 | | |
| *Caucasian* | *N*=1 | *N*=2 | | |
| *Other* | *N*=1 | *N*=0 | | |
| ***BDI Progression*** | | | | |
| BDI at Session 1 | 4.63 ± 4.9 | 24.7 ± 8.25 | -6.3 | **<0.001** |
| BDI at Session 2 | 4.5 ± 4.1 | 15.25 ± 11 | -2.6 | **0.014** |
| *Remitted* | | *N*=7 | | |
| BDI at Session 3 | | 19.2 ± 13.2 | | |
| *Remitted* | | *N*=4 | | |
| **Clinical Variables** | | | | |
| *Combat Trauma* | *N*=5 | *N*=0 | | |
| *mTBI* | *N*=5 | *N*=5 | | |

**Supplementary Table 2: Demographic, clinical, and psychological variables of the test-retest subjects.** (Abbreviations: MDD, Major Depressive Disorder; non-MDD, subjects with no current or prior diagnosis of MDD; F, female; M, male; BDI, Beck Depression Inventory-II score; mTBI, mild traumatic brain injury.)

| | *Cluster 1* | *Cluster 2* | *Unclassified* | $\chi^2$/F | *p-value* |
|---|---|---|---|---|---|
| *Females* | 16 | 18 | 16 | 1.03 | 0.60 |
| *Males* | 39 | 29 | 29 | | |
| *High Pain Average* | 0 | 47 | 28 | 98.8 | **<0.0001** |
| *Low Pain Average* | 55 | 0 | 17 | | |
| *Healthy Controls* | 24 | 21 | 12 | 4.02 | 0.13 |
| *Mixed Psychiatric* | 31 | 26 | 33 | | |
| *Mean BDI* | 9.9 ± 11.7 | 12.6 ± 14.2 | 13 ± 8.4 | 0.93 | 0.40 |
| *Mean Age (years)* | 27 ± 7.6 | 29.9 ±6.2 | 27.2 ± 6.4 | 2.78 | 0.07 |
| ***Predictive ROIs*** | | | | | |
| *Ins. Posterior Long L* | 15 | 12 | 10 | 0.34 | 0.84 |
| *Ins. Posterior Long R* | 18 | 13 | 14 | 0.31 | 0.85 |
| *Ins. Anterior Short L* | 18 | 24 | 17 | 3.70 | 0.16 |
| *Ins. Anterior Short R* | 17 | 23 | 21 | 4.11 | 0.13 |
| *Ins. Middle Short L* | 15 | 21 | 21 | 4.94 | 0.08 |
| *Ins. Middle Short R* | 14 | 21 | 9 | 7.52 | 0.02 |
| *Ins. Posterior Short L* | 17 | 16 | 18 | 0.92 | 0.63 |
| *Ins. Posterior Short R* | 16 | 17 | 19 | 1.89 | 0.39 |
| *Ins. Anterior Inferior Cortex L* | 14 | 14 | 12 | 0.25 | 0.88 |
| *Ins. Anterior Inferior Cortex R* | 13 | 13 | 11 | 0.24 | 0.89 |
| *Ins. Anterior Long L* | 14 | 10 | 16 | 2.50 | 0.29 |
| *Ins. Anterior Long R* | 17 | 17 | 6 | 6.66 | 0.04 |
| *Caudate Nucleus L* | 17 | 17 | 13 | 0.61 | 0.74 |
| *Caudate Nucleus R* | 16 | 17 | 9 | 2.96 | 0.23 |
| *Nucleus Accumbens L* | 25 | 18 | 16 | 1.11 | 0.58 |
| *Nucleus Accumbens R* | 25 | 19 | 15 | 1.52 | 0.47 |
| *Putamen L* | 13 | 13 | 5 | 4.13 | 0.13 |
| *Putamen R* | 10 | 10 | 8 | 0.23 | 0.89 |
| *Pallidum L* | 17 | 16 | 10 | 1.67 | 0.43 |
| *Pallidum R* | 15 | 18 | 11 | 2.40 | 0.30 |
| *Substantia Nigra L* | 23 | 16 | 15 | 0.98 | 0.61 |
| *Substantia Nigra R* | 16 | 17 | 18 | 1.37 | 0.50 |
| *Anterior Cingulate L* | 12 | 14 | 9 | 1.41 | 0.49 |
| *Anterior Cingulate R* | 17 | 8 | 8 | 3.62 | 0.16 |
| *Amygdala L* | 22 | 24 | 10 | 8.24 | 0.02 |
| *Amygdala R* | 22 | 15 | 18 | 0.89 | 0.64 |

**Supplementary Table 3: Cluster classification breakdown.** Subjects are separated based on mixAK cluster classification. Predictive ROIs indicate the number of models (or subjects) which included a given region (e.g., the insula posterior long gyrus on the left side, etc.) within each group. (Abbreviations: BDI, Beck Depression Inventory-II score; L, left cortical side; R, right cortical side.)

### Supplementary Text

#### Supplementary Material 1: Detailed Explanation of Pain-Anticipation Paradigm

Each fMRI session included two acquisitions (i.e,, Run 1 and Run 2) of the pain-anticipation paradigm, separated by 5-7 mins. For the purposes of the aforementioned study performed by Wager et al., 2013, the levels of heat used to invoke painful and non-painful responses were based on post-stimulus self-report [28]. Thus, the temperatures reported as painful and non-painful differed across subjects, and the same temperature could be rated as both painful in one trial and non-painful in another. To avoid any variation introduced by subjective pain ratings, the current study uses two predetermined and consistent temperatures across subjects. The stimulation was delivered through a 9cm$^2$ thermode (Medoc TSA-II, Ramat-Yishai, Israel) on the participant's left forearm, as described elsewhere [19]. The schedule of stimuli differed between imaging runs in a pseudorandom and counterbalanced order. The periods of anticipation were ten seconds long and began with a cue that signaled high, low, or uncertain level of pain. More specifically, ten seconds prior to the onset of pain, the participants were presented with an image of a colored cross. A red cross indicated a temperature stimulus producing moderate levels of heat pain or "high-pain, HP", a green cross indicated a temperature stimulus producing low levels of heat pain or "low-pain, LP", and a yellow cross, indicated pain of uncertain intensity (at 50% probability being high or low, which was not known to the subject). These anticipation periods were followed by seven seconds of either high or low-pain. The high-pain stimulation was administered at 47.5ºC and the low-pain stimulation at 45.5ºC, both of which had a rise and fall rate of 10ºC/sec. Note that both levels of temperature simulations were painful to the subject and they were not informed that only two levels of temperature stimulation would be delivered. In all instances when the level of pain was cued, the participant received the corresponding level of pain. When an uncertain cue was given, the participant was administered either the HP or LP stimulation. Each temperature stimulus was followed by a period of rest, signaled by a change in the color of the cross to blue, that was jittered between 24 to 30 seconds (aside from the short period of rest before the first anticipation cue in each session, which lasted 7 and 10 seconds, respectively). Each session included 14 separate anticipation-pain conditions and lasted a total 618 seconds. In Run 1, there were three HP-cued conditions and four LP-cued conditions. The other seven conditions began with an uncertain cue (UN), three of which were followed by low-pain delivery, and four with high-pain. In Run 2, there were four HP conditions, three LP conditions, and of the seven UN conditions, four were followed with low-pain and three were followed with high-pain. In combination, there was a total of seven HP, seven LP, and fourteen UN (with seven LP and seven HP) conditions (**Supplementary Figure 1**).

**Supplementary Material 2: CONN Preprocessing Pipeline.**

Preprocessing was completed using the default preprocessing pipeline in CONN. The pipeline included the following consecutive steps: (1) functional realignment and unwarp, (2) functional center to (0,0,0) coordinates, (3) functional slice-timing correction, (4) functional outlier detection, (5) functional direct segmentation and normalization, and (6) functional smoothing. For the functional outlier detection (step 4) the intermediate settings were chosen with 97$^{th}$ percentile in the normative sample. For segmentation and normalization (step 5), default tissue probability maps were used for the simultaneous segmentation of gray, white and cerebrospinal fluid (CSF) and Montreal Neurological Institute (MNI) coordinate normalization. The smoothing kernel used in the functional smoothing (step 6) was 4mm full-width halfmaximum (FWHM). Next, denoising was performed on the functional data. For denoising, linear detrending and regression of the confounding effects of realignment and scrubbing was completed. Despiking was implemented before regression and a band pass filter of [0.008Hz, infinity] was applied after regression.

**Supplementary Material 3: Full Cohort Demographics.**

Within the test group eleven females had previously recovered from anorexia (RAN), thirty-five (17 females) met criteria only for major depressive disorder (MDD), nineteen males experienced trauma and had incurred a mild traumatic brain injury (mTBI) during combat, and twenty-five males suffered from combat-related post-trauma stress disorder (PTSD), MDD, and/or mTBI (see Supplementary Material 3). There were no significant differences in age, race, marital status, or gender distribution between healthy and test groups. There was a statistically significant difference in BDI scores between the two groups ($t=-9.75$, $p<0.001$, $df=106$). Healthy controls had an average BDI of $1.9 \pm 3.1$ which falls in the "no depression" range. The test group had an average BDI of $17.9 \pm 12$ which indicates the presence of mild or moderate depressive symptoms. Of the thirty-two test-retest subjects, twenty-four (7 females) met criteria for MDD at the first imaging session. The demographic and behavioral attributes of the test-retest subject are outlined in Supplementary Material 4.

**Supplementary Material 4: Test-Retest Subjects.**

Within the test-retest cohort, a BDI-2 score was obtained for each subject at each session. The average BDI-2 score for both groups, along with the number of subjects who entered remission or sub-MDD classification at either session 2 or session 3, is included. At the second session, twelve of the original twenty-four MDD subjects still met criteria for MDD, seven had gone into remission and three were considered sub-MDD. Of the eleven subjects who completed a third scanning session, all were diagnosed with MDD at Session 1; three had remitted by Session 2 and remained in remission at Session 3, one remitted between Session 2 and Session 3, and seven still met criteria for MDD at Session 3. BDI-2 scores between sessions reflected these changes in MDD diagnosis. There was not a significant difference between the two stable groups and remittance rates (chi = 1.22, p = 0.54), or between the stable and unstable groups and remittance rates (1.37, p = 0.50).

**Supplementary Material 5: Optimal LASSO model parameters.**

Optimal parameters for the LASSO model were: alpha optimized based on single-subject accuracy, lambda optimized based on the minimum cross-validated lambda per subject over one hundred validation runs, and misclassification error ("class") used as the loss parameter. In accordance with previous findings [47], the t-statistic was found to be superior to the beta-coefficient-based activation maps. Model performance is depicted in **Supplementary Figure 2**.

**Supplementary Material 6: Most Deterministic Regions**

The adaptable single-subject LASSO model was designed to select the regional neural activities within 26 ROIs most significant to distinguish between certain anticipations of high and low-pain intensity. On average, 8.1 ± 2.8 regions were selected as variables with predictive value in single-subject LASSO models. Frequency map of regions of interest included in single-subject LASSO is depicted in **Supplementary Figure 3**. The most frequently included regional neural activity predictor of low versus high-pain anticipation were Nucleus Accumbens and anterior short insular gyrus on the right hemisphere (**Supplementary Figure 3**). Activation within these regions were included in 94 (64%) and 93 (63%) subjects' LASSO models, respectively. At a significance level of 0.05 with 26 regions, significance was determined at $p<0.019$. There was no significant difference in incidence of regional determinism between healthy controls as compared to the "mixed psychiatric" test group. Within test-retest subjects the same regions were highly deterministic.

**Supplementary Material 7: LASSO Model Stability**

The models were consistent in the selection of relevant predictors across runs and between cohorts; this stability is illustrated by the similarity matrices in **Supplementary Figure 4**. These matrices also show the immense variability in LASSO-selected predictors between subjects, further supporting the use of single-subject analysis. Stability of the LASSO model was also seen between analyses of separate cohorts. Since the test-retest cohort was comprised of a subset of the full cohort, the corresponding models were compared to assess LASSO stability across analyses. The inset image (top left, **Supplementary Figure 4**) represents the consistency between the model created for subject #52 during full cohort analyses and during test-retest analyses. Of the thirty-two test-retest subjects, 29 (90.6%) of the subjects' models included the same predictors as the model created in the full cohort processing.

**Supplementary Material 8: Clinical Subgroups vs. Cluster Classification.** Histograms depicting average prediction of subjects versus cluster classification within the full cohort (**Supplementary Figure 5a**), and subgroups of healthy controls (**Supplementary Figure 5b**), MDD (**Supplementary Figure 5c**), RAN (**Supplementary Figure 5d),** PTSD/mTBI with combat trauma (**Supplementary Figure 5e**), MDD/mTBI (**Supplementary Figure 5f**). There was no significant difference in cluster classification within any of the aforementioned groups. Subjects clustered only based on average predictions such that all subjects in cluster 1 had average predictions greater than 0.5, cluster 2 had average predictions less than 0.5, and unclassified subjects had predictions between 0.25-0.75.
